# Mode and tempo of microsatellite length change in a malaria parasite mutation accumulation experiment

**DOI:** 10.1101/560516

**Authors:** Marina McDew-White, Xue Li, Standwell C. Nkhoma, Shalini Nair, Ian Cheeseman, Tim J.C. Anderson

## Abstract

Microsatellite sequences are widely assumed to evolve neutrally, but also play an important role in bacterial pathogenesis, human disease and transcript abundance. The malaria parasite *Plasmodium falciparum* genome is extraordinarily AT-rich, containing 132,449 microsatellites-stretches of perfect 1-9 bp repeats between 10-1000bp, which comprise 10.74% of the 23 Mb genome. This project was designed to determine the mode and tempo of microsatellite mutations in malaria parasites. We maintained 31 parasite lines derived from a single 3D7 parasite cell for 114-267 days, with frequent bottlenecking to a single cell to minimize effective population size, allowing us to measure mutations accumulated over ~13,207 mitotic divisions. We Illumina sequenced the genomes of both progenitor and end-point mutation accumulation (MA) parasite lines in duplicate to validate stringent calling parameters. Calls were 99.89% (GATK), 99.99% (freeBayes) and 99.96% (HipSTR) concordant in duplicate sequence runs from independent sequence libraries. We observed 98 microsatellite mutations, giving rates of 2.11 × 10^-7^ - 1.46 × 10^-8^ /cell division that were strongly influenced by repeat motif and array length. Mutation rate was low relative to other organisms. However, despite this, in a single infection (10^11^ parasites) there will be 1.46 × 10^3^ - 2.11 × 10^4^ independent mutations at any single microsatellite locus. Given that many microsatellites are found in promotors, introns, within or close to coding sequences, we suggest that they may be important regulators of transcriptional and phenotypic variation in this pathogen.

**Author summary:** Mutation is central to evolution: in pathogens, the rate of mutation may determine how rapidly drug resistance evolves or how effectively pathogens can escape immune attack. Malaria parasites have small extremely AT-rich genomes, and genetic variation in natural populations is dominated by repeat number changes in short tandem repeats (microsatellites) rather than point mutations. We therefore focused on quantifying microsatellite mutation. We established 31 parasite cultures in the laboratory all derived from a single parasite cell. These were maintained for 114-267 days with frequent reductions to a single cell, so parasites accumulated mutations during ~13,207 cell divisions. We sequenced the parasite genomes at the end of the experiment to count the mutations. We highlight several conclusions: like other organisms studied, microsatellite mutation rates are associated with both repeat number and repeat motif. However, 41% of changes resulted from loss or gain of more than one repeat: this was particularly true for long repeat arrays. Unlike other eukaryotes, we found no insertions or deletions that were not associated with repeats or homology regions. Overall, we found that microsatellite mutation rates in malaria were amongst the lowest recorded and comparable to those in another AT-rich protozoan (the slime mold *Dictyostelium*).

## Introduction

The genome of *Plasmodium falciparum* malaria parasites is 80% AT rich. As a consequence, microsatellite repeats, defined here as strings of repeated motifs 1-9bp, are extraordinarily common in malaria parasite genome. There are 132,449 microsatellites in the 3D7 reference genome (PlasmoDB,http://plasmodb.org/common/downloads/release-32/Pfalciparum3D7/): this equates to one microsatellite every 173bp and 10.74% of the 23 Mb genome is composed of microsatellites [1]. These markers typically have multiple alleles and high heterozygosity in natural populations making them the dominant source of genomic variation in malaria parasites [2]. In contrast, microsatellites constitute just 1-3% of the genome in organisms such as *Drosophila* or humans with AT/GC content closer to 50% [3, 4].

Microsatellites are generally assumed to evolve neutrally, and have been widely used as genetic markers for population genetic and epidemiological studies, including for malaria parasites [5]. However, the assumption of neutrality is questionable. Repeat mutation within promotors, introns, or coding regions can alter transcription through interference with transcription factor binding, changing spacing between regulatory elements, or by their effects on alternative splicing [6]. It has been suggested that microsatellites may act as evolutionary capacitors fine-tuning and optimizing transcript levels. The effects on transcription may be extensive: 10-15% of heritability in human gene expression is estimated to result from associated cis-acting microsatellite loci [7]. Microsatellite repeat expansions are involved in at least 40 different human diseases, including fragile X syndrome or Freidrich’s ataxia [8].

In many bacterial pathogens, loss or gain of repeats within promotors or coding sequences are involved in “phase shifts” resulting in changes in pathogenesis. For example, *Campylobacter jejuni* has 29 phase variable (or contingency loci) that are stochastically switched on or off by loss or gain of monomeric G/C repeats, which put genes in or out of frame [9]. In fact, repeat tracts are used for identifying genes involved in pathogenesis in bacteria [10, 11]. Pathogens typically go through extreme bottlenecks during transmission from one host to another. In the case of bacteria, rapid mutation of phase variable loci may provide a mechanism for stochastic generation of phenotypic variation from a few colonizing bacteria. Similarly, a single malaria sporozoite inoculated from an infected mosquito can produce a human infection containing up to 10^11^ blood stage parasites. Rapid mutation of microsatellites that influence transcription has the potential to generate extensive phenotypic variation on which selection can act in blood stage malaria infections.

The extremely large number of repeat loci in malaria parasites, and their potential impact on phenotypic variation, make studying microsatellite evolution a priority. Scoring repeat variation using next generation sequence data is problematic for several reasons: (i) current Illumina read length is similar to length of microsatellite arrays, which imposes an upper limit on what loci can be effectively scored (ii) amplification steps during library preparation typically generate “stutter”, generating a distribution of repeat array sizes which complicates scoring. Two groups have examined mutation rate in *Plasmodium*. Bopp et al [12] examined SNP mutation while a separate experiment examined both SNP and indel mutations [13, 14]. While these studies inferred mutation rates from clone trees, we aimed to use a classical mutation accumulation experimental design in which multiple parasite clones derived from a single progenitor maintained over multiple generations with frequent bottle-necking to a single cell. A central aim of this work was to evaluate robustness of different methods for scoring spontaneous microsatellite or indel polymorphism in AT-rich malaria genomes in a mutation accumulation experiment. We then aimed to evaluate the impact of repeat motif, array length and genomic context, gene function, and transcriptional activity on observed mutation rates.

## Results

### Mutation accumulation lines

To investigate the mutation rate in the *P. falciparum* genome, we generated long-term mutation-accumulation (MA) lines starting with a randomly chosen single clone from laboratory adapted 3D7 (Fig 1) generated by dilution cloning. The MA lines passed through a single-cell bottleneck every 21(±4) days to minimize the strength of selection and to fix mutations within each line. Our MA experiment allowed the accumulation of mutations over an average of 170 (range: 114-267) days in 31 independent MA lines (S1 Table). This equates to a total of 85 (57-134) 48-hr asexual generations or 425 (285-668) mitotic divisions: in total 2,641.5 48-hr asexual generations, or an estimated 13,207 mitotic divisions.

**Fig 1.**
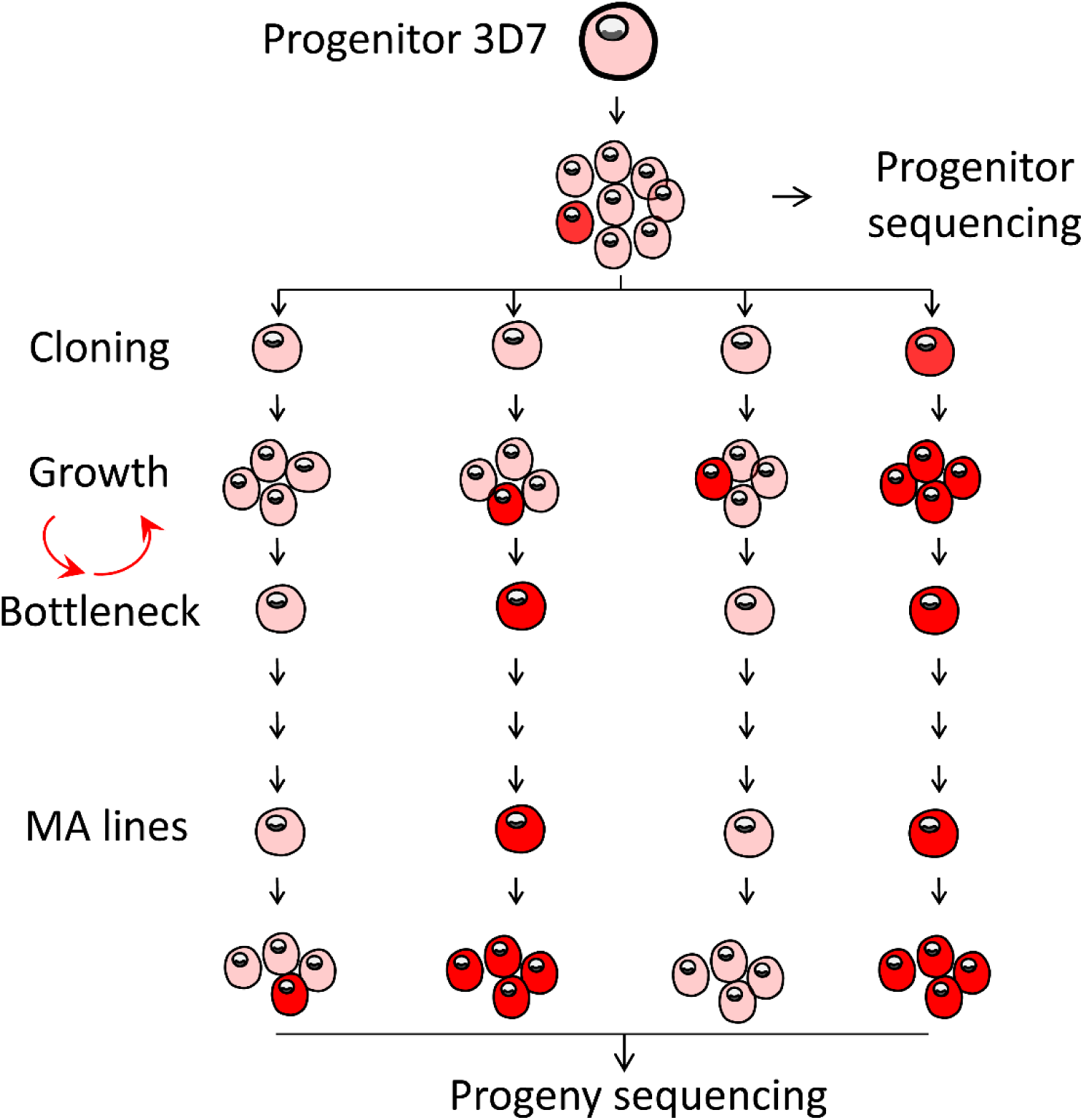
Generating of mutation accumulation lines with *P. falciparum*. The progenitor 3D7 parasite was dilution cloned prior to the experiment, expanded, and then dilution cloned again to found each MA line, with additional cloning steps at frequent (17-25 day) intervals. Transitions from pink to red shading indicate accumulation of mutations.

### Compositions of short tandem repeats in the *P. falciparum* genome

We identified 132,449 perfect microsatellites from the *P. falciparum* 3D7 genome (http://plasmodb.org/common/downloads/release-32/Pfalciparum3D7/) with at least three repeats, 10-1000bp in length and 1-9bp per repeat unit, which accounts for 10.74% of the whole genome. Among these microsatellites, 123,834 (93.50%) of them are located in the core genome (defined in [2]). As genotyping microsatellites that exceed 70bp in size invariably requires reads longer than 100bp [15], and the genotype call rate declined for longer microsatellites, we restricted this analysis to microsatellites with a size range 10-70bp. This accounts for 99.9% (123,722/123,834) of the core genome microsatellites and provides a good representation of microsatellites from the complete *P. falciparum* genome (S1 Fig and S2 Table).

Microsatellites with a 1-6bp repeat unit account for 98.16% of all the *P. falciparum* microsatellites. Of these, 37.68% (46,626 of 123,722) are homopolymeric tracts, and 35.48% (46,849/123,722) are dinucleotide repeats (S1 Fig). 94.64% of the microsatellites have a repeat unit containing A and T: this includes 99.94% (46,600 of 46,626) of homopolymeric tracts, 99.05% (43,490 of 43,903) of dinucleotides, and 82.38% (10,973 of 13,106) of trinucleotides.

Microsatellites are abundant in both coding and non-coding regions (S2 Fig). The total number of microsatellites located in coding DNA sequences (CDS), intergenic, intronic and promoter (within 1kb upstream of the initiation codon ATG) regions are 19,912 (16.09%), 35,575 (28.75%), 19,086 (15.43%) and 49,149 (39.73%), respectfully. Microsatellite densities vary in different sequence categories. We observed 10.35, 13.56 and 11.28 microsatellites per kb in intergenic, intronic and promoter regions, but only 1.72 microsatellites per kb in the CDS region (S3 Table). These distributions are significantly different from random expectations (*p* < 2.2×10^-16^, Pearson’s Chi-squared test). The average repeat numbers for microsatellites in CDS region were also significantly lower than those located in non-coding regions (*p* < 2.2×10^-16^, two sample Wilcoxon rank sum test), with average repeat numbers of 12.0, 11.9 and 11.9 in intergenic, intronic and promoter regions respectively but only 8.2 in CDS region (S2 Fig).

The repeat motifs of microsatellites were highly biased in both coding and non-coding regions (S2 Fig and S3 Table). More than 78.06% of trinucleotides are located in the CDS region while dinucleotides (1.89%), tetranucleotides (2.49%) or pentanucleotides (3.68%) are very rarely located in the CDS. The density of trinucleotides was still 2.72-3.73 times higher in CDS region than in all the other non-coding regions while the densities of dinucleotides, tetranucleotides and pentanucleotides in non-coding regions were 50.40-77.13 times greater than those in the coding regions.

We also examined *P. falciparum* minisatellites (defined as repeats 30-1000bp in length). We identified 3,528 minisatellites from the 3D7 genome with at least three repeats, and 10-30bp per repeat unit, of which 3,294 (93.37%) are located in the core genome. 61.35% of minisatelllites are located in gene coding region (S3 Fig and S4 Table), which is much more than expected by chance (*p* = 1.19×10^-5^, Pearson’s Chi-squared test). Unlike microsatellites, the repeat number of minisatellites located in coding and non-coding regions is not significantly different (*p* = 0.035, two sample Wilcoxon rank sum test). Only 58.08% (1,913) of the minisatelllites are smaller than 70bp, and therefore detectable by short read Illumina sequencing.

### Concordance of genotype calling in duplicated runs

We made two independent sequencing libraries for both the 3D7 progenitor and each of the 31 MA lines, with an average coverage depth of 91.8X and 2,141 Mb 101-bp paired-end reads sequenced for each library (S1 Table). To reduce false positives due to alignment errors, we excluded highly variable genome regions (subtelomeric repeats, hypervariable regions and centromeres) and only performed genotype calling in the 20,782 kb core genome (defined in [2]). We called genotypes using three methods (GATK, freeBayes and HipSTR). We filtered the initial mutation call set from each of the three methods using two approaches: (i) a standard method (using recommended filters) and (ii) a stringent method, using optimized filtering parameters (Fig 2). We then computed genotype concordance for each replicate pair as (1-[number of variants with a discordant genotype call]/[total number of variants with non-missing genotype calls in both sequence runs]).

**Fig 2.**
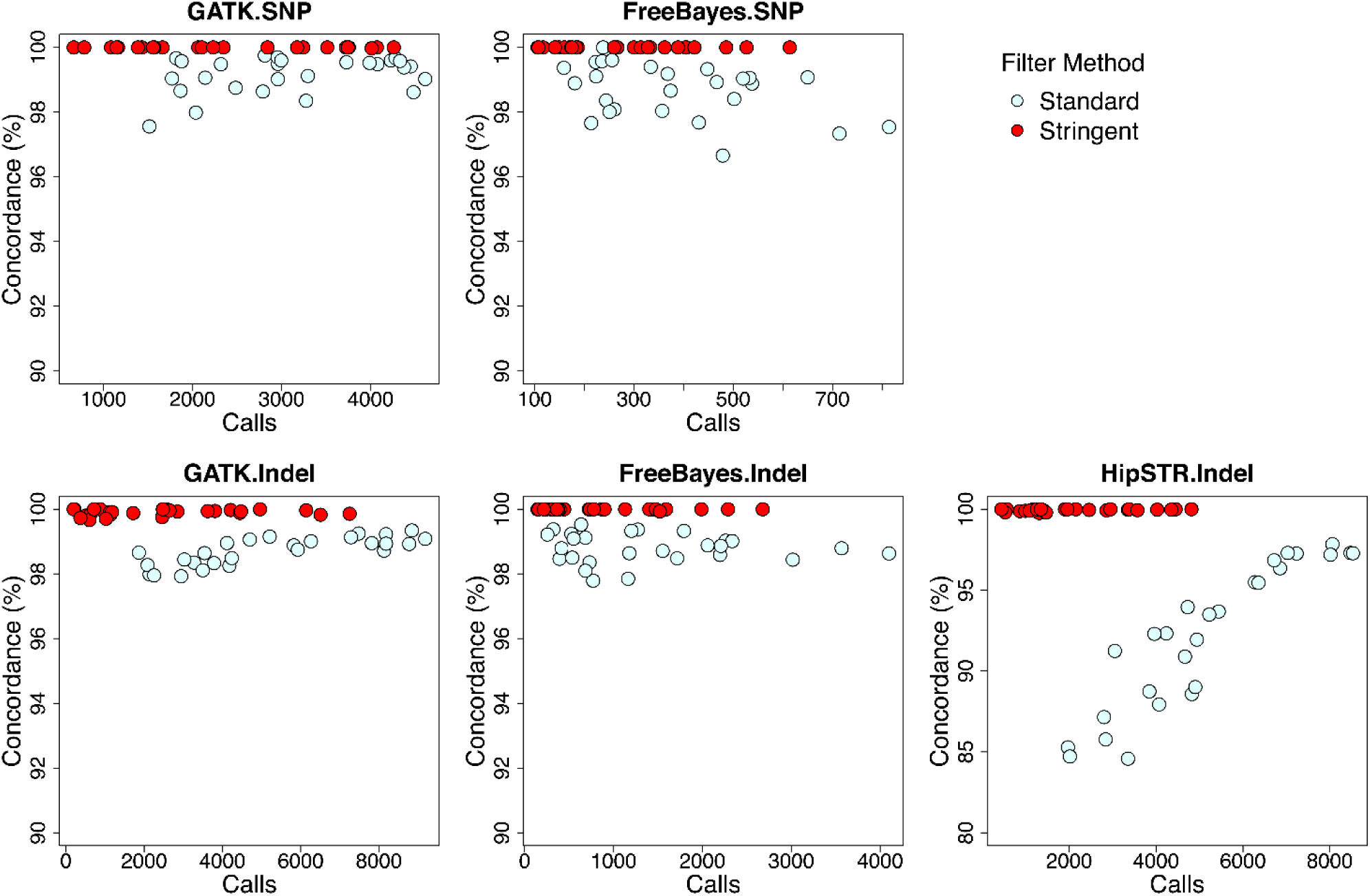
Comparison of different genotyping and filter methods. See text for parameters used for standard versus stringent filtering. Stringent filtering allowed high concordance calling in duplicate sequence runs for scoring both SNP and indels.

Under the standard filtering parameters, the average concordances were 99.16% (GATK) and 98.72% (freeBayes) for SNP callings, and 98.70% (GATK), 98.80% (freeBayes) and 92.22% (HipSTR) for indel or microsatellite calls. These concordance rates are sufficiently accurate for scoring variation in populations or genetics crosses, but are insufficiently accurate for scoring rare mutation in MA lines, because there will be a high proportion of false positives.

We therefore implemented more stringent filter methods (S5 Table). We first explored the performance of different filters for optimizing genotyping calls. Two filtering parameters−the percentage of supporting reads to current genotype (purity) and the likelihood ratio for the called genotyped relative to the second best genotype (PLdiff) −were particularly effective for minimizing discordant genotype calls (Fig 2, S4 Fig). The concordance rates were dramatically improved, reaching 100% (GATK) and 100% (freeBayes) for SNP callings, and 99.90% (GATK), 99.99% (freeBayes) and 99.95% (HipSTR) for indel or microsatellite calling (Fig 2): sufficiently accurate for rare mutations discovery in this study.

We further removed the remaining false positives by visual inspection of all the putative mutations detected (S5 Fig). We retained 13/20 (65.0%) of SNPs identified from stringently filtered GATK calls and 13/15 (86.7%) of SNPs identified from stringently filtered freeBayes calls. We retained 35/45 (77.8%) indels identified from our stringently filtered HipSTR calls set following visual inspection. Indel calls from GATK and freeBayes were retained at a higher rate than those from HipSTR (81/99 [81.8%] for GATK and 43/50 [86.0%] for freeBayes) after visual inspection (S6 Table).

### Mutations observed in mutation accumulation experiment

We included 17 SNPs and 106 indels from 31 MA lines in the mutation rate analysis (Fig 3). For the 17 remaining SNPs: 9 were detected by both freeBayes and GATK, while 4 were detected by GATK only and 4 by freeBayes only (S7 Table). We detected 106 indels in the mutation accumulation analysis. These indels included 81/106 (76.4 %) identified by GATK, 40.6 % by freeBayes, 33.0% identified by HipSTR. Just 11/106 (10.4 %) were identified using all three methods (S8 Table).

**Fig 3.**
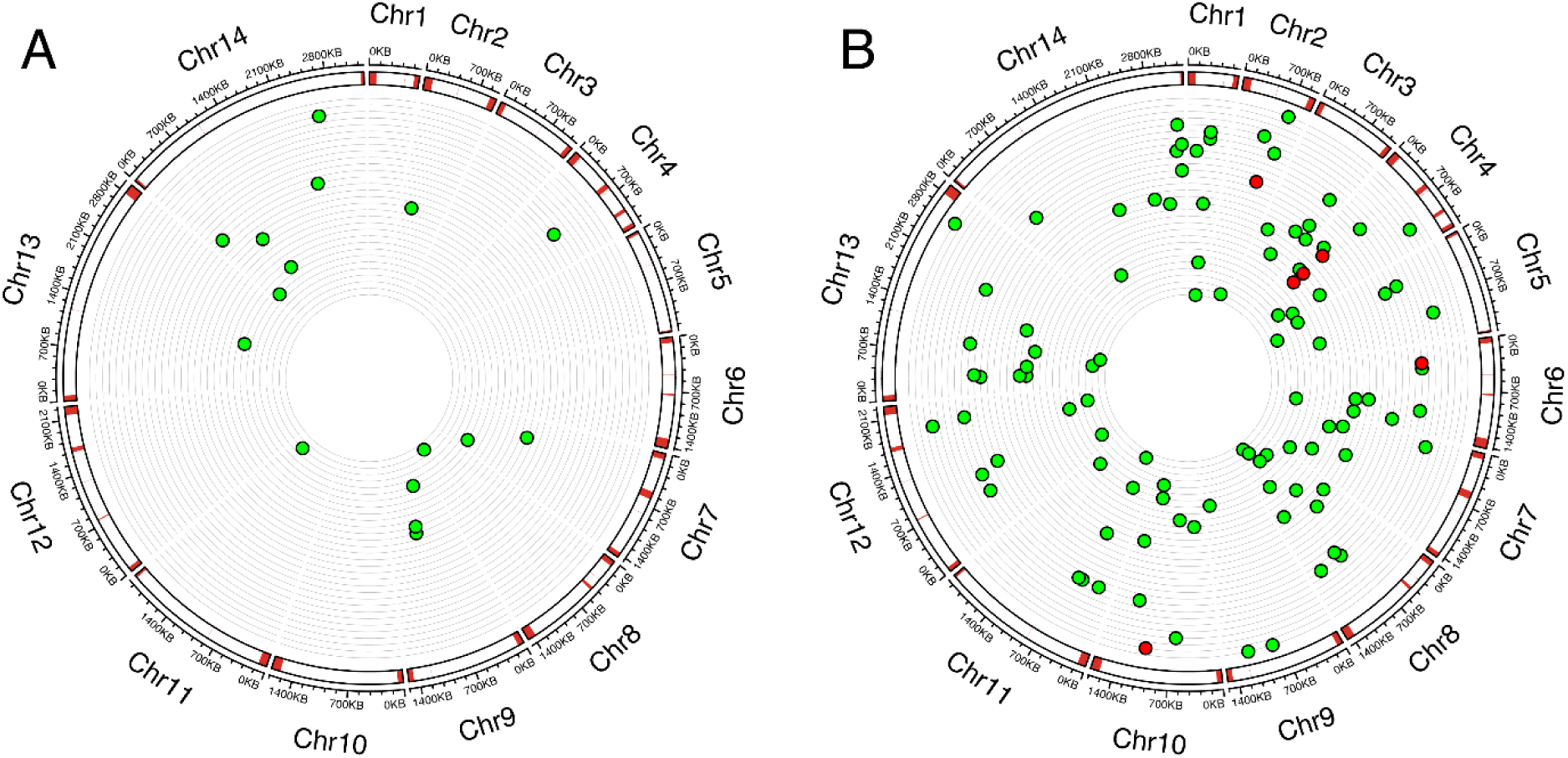
The genome location of (A) base-substitutions and (B) indels in the 31 independent MA lines. From the outer to inner circles: kilo base scale of 14 chromosomes; location of variable regions (subtelomeric repeats, hypervariable regions, and centromeres) of the *P. falciparum* genome are marked in red; position of each microsatellite mutation or base-substitution in MA lines, with each ring representing the genome of an individual MA line.

### Base substitutions

There were 17 base substitutions in 31 MA lines maintained for cumulative total of 5,283 culturing days (Fig 3A and S7 Table), giving an estimated mutation rate to be 3.10±0.75 × 10^-10^ base substitutions per site per asexual cycle.

We analyzed mutation spectra in the MA lines by considering the six nonstrand-specific base-substitution types (Fig 4A). The Ts/Tv ratio (0.31) is not significantly different from the random expectation of 0.5, but the transition and transversion subtypes occurred at different rates. G:C->A:T transitions had higher mutation rates, while A:T->G:C transitions and A:T->C:G transversions had lower rates compared to expectation (*p* < 0.05 for all comparisons, two-sided exact binomial test).

**Fig 4.**
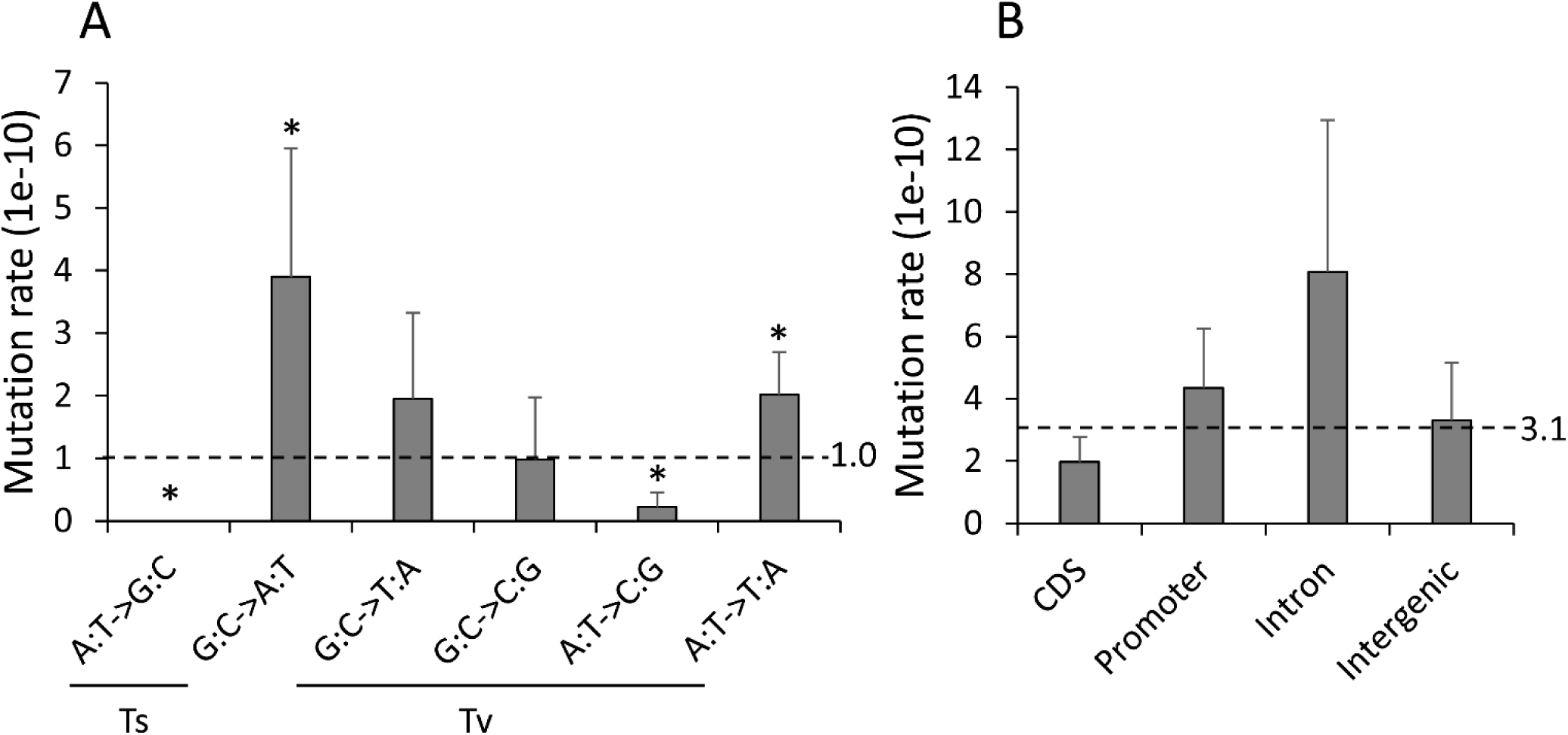
Base substitutions rate. (A) Mutation spectrum across six base-substitution types. (B) Mutation rate for different coding sequence categories. * indicates *P* value < 0.05.

We compared the distributions of mutations in coding sequence categories: CDS, promoter, intron, and intergenic regions. Mutation rates were not significantly different in these four functional sequence categories (Fig 4B), suggesting equal mutational susceptibilities for each of these categories in the MA-line genomes and further corroborating that selection has minimally affected the pool of mutations analyzed here. We observed 6 nonsynonymous mutations and 1 synonymous mutations in exon sequences. These numbers were not significantly different (*p* = 1, Pearson’s Chi-squared test) from the expected values based on the nonsynonymous to synonymous codon potential ratio estimated by Bopp et al [12].

### Microsatellites, insertions and deletions

We detected 106 indels in 31 MA lines. These are approximately 6 times as common as base substitutions (S8 Table). 94.3% (100) of indels are found in arrays of tandem repeats: these include 98 microsatellites and 2 minisatellites. All the microsatellites and minisatellites involved loss or gain of complete repeat units. We also detected 6 indels, outside of tandem repeats: all were deletions (average size = 18.3bp, range = 15-26bp) with homology sequences (10-26bp) nearby (Fig S6). We didn’t detect small indels (1-4bp) outside of microsatellite or larger indels that contained apparently random DNA sequences (S8 Table). We found a similar pattern to the 3D7 MA lines analyzed by Hamilton et al [14], who observed 156 microsatellite indels, 7 minisatellite indels and 1 non-tandem repeat (with homology indel) among 164 indels. In comparison, 69.9% indels in humans are not associated with repeat sequences [16].

We analyzed the genomic distribution of the indels. 2/6 non-variable number tandem repeat (non-VNTR) indels and all (2/2) minisatellite indels are located in the coding region, while only 15/98 microsatellite changes are observed in coding region. We found a similar distribution of indels to Hamilton et al [14]: they observed 1/1 non-VNTR indels, 7/7 minisatellite indels and 8/156 microsatellites are located in coding region. The distribution of microsatellite indels is significantly different from minisatellite indels (*p* = 2.91×10^-8^, Pearson’s Chi-squared test), which may be due to restriction of microsatellite in coding region: 2,021/3,294 (61.35%) of the minisatelllites, but only 19,912/123,722 (16.09%) of the microsatellites are located in gene coding region. All (15/15) of the microsatellites indels in the coding region are divisible by 3 (Table S8). The non-coding region (6/83) contains fewer repeats evenly divisible by 3 compared to the coding region (*p* = 7.54×10^-7^, Pearson’s Chi-squared test).

### Mode and tempo of microsatellite mutations

We found 98 microsatellite mutations in our MA experiment. The overall mutation rate for *Plasmodium* microsatellite is 3.00±0.30 × 10^-7^ per locus per asexual cycle, which ~ 1000 times higher than base substitution rate (3.10±0.75 × 10^-10^ base substitutions per site per asexual cycle). Dinucleotide mutations are the most common microsatellite mutation (Fig 5). As mutations for microsatellites bearing repeat motif size >6 are rarely detected, we compared mutation rate for microsatellites with homonucleotide to microsatellites with hexanucleotide repeat motifs (n=96) (Fig 5C). The mutation rate increased as motif size increased. Pentanucleotides showed higher mutation rate compare to expectations (*p* = 7.36×10^-3^, two-sided exact binomial test), while the homopolymer mutation rate was lower than expected (*p* = 2.60×10^-10^, two-sided exact binomial test). We detected increased mutation rates as microsatellite repeat number increased (Fig 5E). The increasing trend is particularly obvious for dinucleotides (Fig 5F): mutation rate increases > 20 fold as repeat number increases from 5-10 to 20-25, and fit to a one-parameter exponential model (S7 Fig, y = 0.27e^0.20x^, *R²* = 0.98).

**Fig 5.**
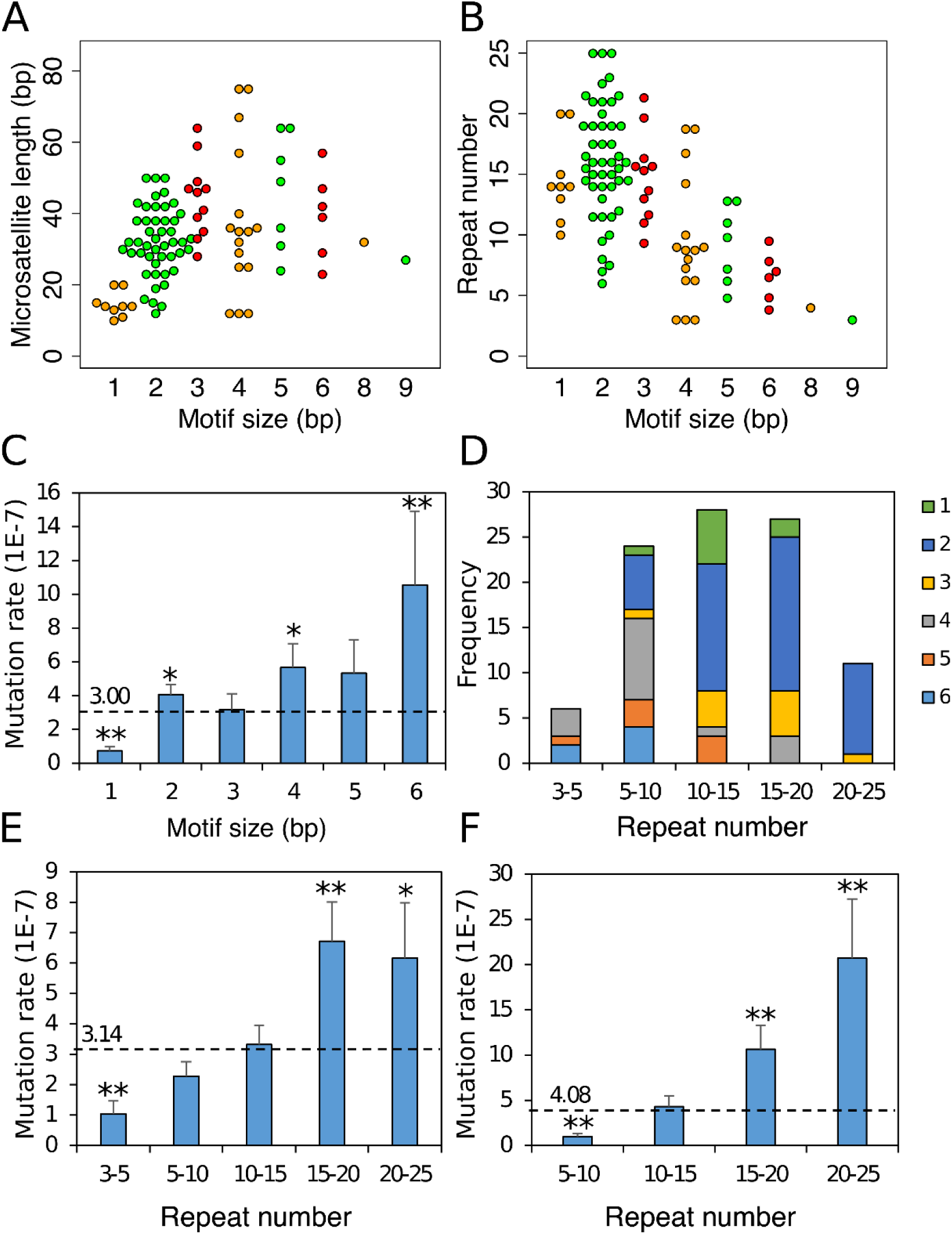
Microsatellite mutation distributions. Spectrum of microsatellite mutations with different motif size, length (A) and repeat number (B). (C) Mutation rate for different motif sizes. (D) Composition of mutations by repeat number and motif size. (E) Mutation rate at different repeat number. (F) Dinucleotide mutation rate at different repeat number. * indicates *P* value < 0.05; ** indicates *p* value < 0.01.

The distribution of microsatellite sequences varies among genomic regions (Fig 6). There are no mononucleotide, dinucleotide, tetranucleotide or pentanucleotide mutations detected in coding regions, and no trinucleotide mutation in noncoding region. There is no significant difference between genomic distribution of base substitutions and microsatellite mutations (*p* = 0.339, df =3, Pearson’s Chi-squared test). The overall numbers in each sequence category do not differ significantly from expectation (*p* = 0.076, df =3, Pearson’s Chi-squared test). The mutation rate in introns is significantly higher than the average (Fig 6A, *p* = 2.84×10^-3^, two-sided exact binomial test).

**Fig 6.**
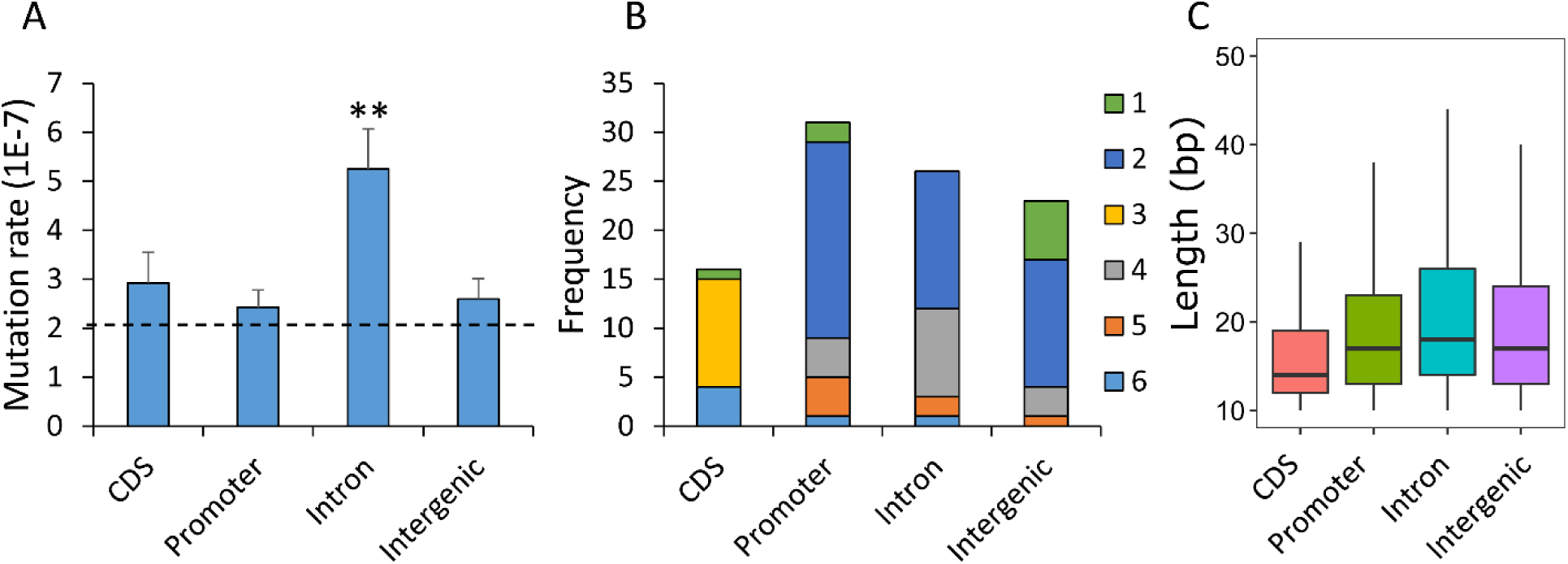
Microsatellite mutation and genomic location. (A) Microsatellite mutation rate. (B) Distribution of mutations for different motifs sizes by genome location. (C) The distribution of microsatellite tracts by genome locations. ** indicates *p* value < 0.01.

Among the 96 microsatellite mutations, there are 43 deletions (1-11 repeat units) and 53 insertions (1-8 repeat units). While most mutations involve loss or gain of single repeat units, the distribution is significantly asymmetric, with 4.5 times more deletions than insertions showing a change of >5 repeat units (Fig 7). We investigated the effect of microsatellite repeat number on the observed bias towards repeat loss using dinucleotide mutations, the most abundant class of microsatellite mutations (Fig 7C and D). The size of insertions and deletions increased dramatically for long microsatellites. We observed mutations averaging 1.75 repeats for microsatellites with ≤ 15 repeats, but averaging 4.07 repeats for microsatellites with > 15 repeats (*p* = 6.99×10^-4^, Wilcoxon rank sum test). Furthermore, the size of deletions was significantly larger than that of insertions (Fig 7D, *p* = 4.30×10^-5^, Wilcoxon rank sum test).

**Fig 7.**
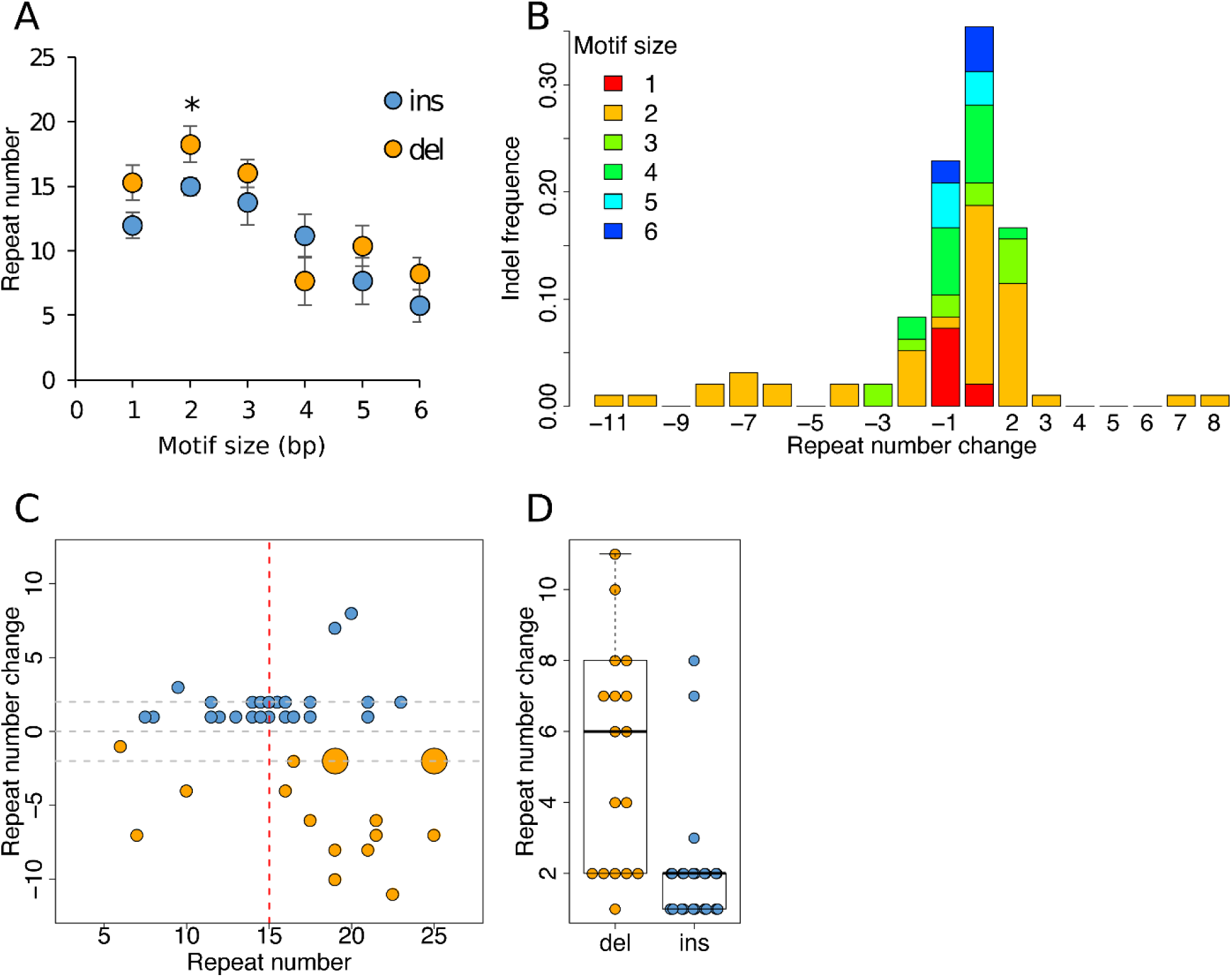
Patterns of insertion and deletion in microsatellites and dependency on array length. (A) Mean repeat number associated with insertions or deletions for different motif sizes. For all motif sizes (except 4bp) insertions are observed in microsatellites with fewer repeats than deletions: this is significant for the most abundant motif type (dinucleotides). (B) Size distribution of insertion and deletion events by motif type. The size distribution of indels is skewed and deletions tend to be larger than insertions. (C) The relationship between indel (+/-) size and repeat number with dinucleotides. Long repeat arrays show larger changes in repeat number than short repeat arrays and there is a significant excess of large repeat losses. Distribution of insertion and deletion size for short (≤15) and long (>15) repeats. (D) Deletions are significantly larger on average than insertions with dinucleotides (see also panel C).

## Discussion

### Scoring methods for indels and microsatellites

Particularly stringent methods are needed for mutation accumulation experiments, where numbers of mutations are expected to be quite modest. We used three different approaches to score microsatellite mutations, with stringent filtering to remove false positives. Our stringent methods filtered calls using purity and PLdiff resulted in 99.63-100% concordance between sequences generated from independent libraries. Even with almost perfect concordance between calls, visual inspection clearly indicated that some of the calls were incorrect. GATK called the most indels with an error rate of 18.2% on manual inspection. FreeBayes and HipSTR called fewer repeat mutations with 14.0 % and 22.2% error rate, respectively. We removed 18% total calls following visual inspection. For comparison Hamilton et al (2017) also removed 16/180 (8.9%) calls for the same reasons [14].

We found that different scoring approaches identify different subsets of mutations from the same dataset. Of 106 indels identified (98 microsatellites and 8 indels), GATK identified the majority (81/106, 76.4%), followed by freeBayes 43/106 (40.6 %) and HipSTR (33.0%). Importantly, only 11/106 calls were made by all three methods: hence we suggest that multiple methods are needed to provide the most complete inventory of indel mutations.

All indel mutations were associated with repeats or homologous regions (S6 Fig). Indels can be generated under multiple mechanisms, such as DNA biosynthesis errors [17], DNA polymerase slippage [18, 19], and repair of DNA double-strand breaks [20]. We speculate that the lack of small random indels in the 3D7 genome relative to humans may be due to the inefficiency of *Plasmodium* end-joining pathway [21].

### Comparison with other *Plasmodium falciparum* mutation rate estimates

<u>Indels and microsatellites</u>: Of the 106 variants identified, 98 were in microsatellites, with just 2 minisatellites and 6 in non-repetitive tracts. This gives an overall microsatellite mutation rate of 3.00 (±0.30) × 10^-7^/asexual cycle. Our results both confirm and extend the analysis by Hamilton et al 2017 [14]. They examined mutation accumulation in 37 sub-lines from a 3D7 clone tree over a total of 203 days in culture (101.5 asexual generations). They found 164 indels, giving an overall microsatellite mutation rate of 4.45(±0.97) × 10^-7^, which is very similar to our estimate. Both our study and Hamilton et al [14] found that the microsatellite mutation rate was higher in introns compared with coding sequence, promotors or intergenic regions. Microsatellites from the intronic regions are longer on average than those from CDS, intergenic and promoter regions (Fig 6C, *P* < 2e-16, ANOVA). When fitting both length and genomic locations into a binomial generalized linear model (GLM), length was a significant positive predictor (Z = 14.959, *P* < 2e-16) for microsatellite mutation, while genomic location had no influence (*P* > 0.05 for all regions). We conclude that the higher intronic mutation rates observed is driven by the longer microsatellites within introns rather than differences in selective constraints among the genome locations.

<u>Single Nucleotide polymorphisms</u> – Our estimate of mutation rates (3.10±0.74 × 10^-10^ base substitutions per site per asexual cycle) are similar to those from two previous experiments that have examined SNP mutation rates in *P. falciparum*. The mutation rate was 5.23 × 10^-10^ from MA lines (3 clones) built by Bopp et al [12], and 2.10 × 10^-10^ from MA lines (37 clones) generated by Hamilton et al [14], with the same calculation equation (see Methods). Bopp et al [12] adjusted the mutation rate by assuming that lethal or deleterious nonsynonymous mutations that arise during long-term culture may not be detected. Their adjusted base substitution rate was 1.7 × 10^-9^ for 3D7 without drug selection. Using that method, our mutation rate becomes 8.14 × 10^-10^ base substitutions per site per asexual cycle. Thus, the 3D7 base substitution rates estimated by different laboratories are highly consistent.

### Mode and tempo of microsatellite mutation

Both the length of repeat arrays and the repeat motif have been shown to impact microsatellite mutation rate in multiple species [22]. Consistent with this, in *P. falciparum*, we see that large microsatellite motifs (4-6bp) show higher mutation rates than motifs <4. There is also a strong relationship between microsatellite heterozygosity and repeat array length [23]. For dinucleotide AT repeats, the dominant repeat class in the *P. falciparum* genome, mutation rate increases from 9.87 × 10^-8^ for 5-10 repeats to 2.07 × 10^-6^ for 20-25 repeats. There is a strong positive relationship between mutation rate and repeat number; this fits better to a one-parameter exponential model (S7 Fig, *y* = 0.27e^0.20*x*^, R² = 0.98, *P* = 0.01; F value = 28.72, *P* = 0.03 compared to the liner model, ANOVA) as observed for di-nucleotide repeats in humans [24].

Stepwise mutation models, in which single repeat units are lost or gained from arrays, are frequently used to model microsatellite evolution. We found similar numbers of losses and gains of repeats (43 deletions vs 53 insertions). This differs from Hamilton et al (2017) [14] who found significantly more insertions than deletions (56 deletions vs 108 insertions). We found several features that fit poorly with a simple stepwise mutation model. Approximately half (59.2%) the mutations involved a single repeat unit while the remainder involved multiple repeat units. Some of these length changes were large: we observed gains of 8 repeats and losses of 11 repeats. Interestingly, we found that mutational size was dependent on array length and highly asymmetric. Long arrays (≥15 repeats) showed larger mutational changes than smaller arrays (<15 repeats). Furthermore, these long arrays also tend to have to have larger deletions than insertions. This asymmetry is expected to limit large repeat expansions [25], such as those responsible for repeat associated disorders in humans [26]. However, it is also possible that shorter read sequencing data limits our ability to score large insertions, resulting underestimation of this class mutations

### Mutation frequency, transcription and gene essentiality

Several studies report an association between mutation rates and proximity to highly expressed genes [27, 28]. One possible interpretation is that mutation results from collisions between replication and transcription machinery, which will occur more frequently in highly expressed genes. To examine the roles of transcription, motif size, array length, genomic context, we fitted all these variables into a binomial family GLM (S9 Table). As expected motif size and array length were significant positive predictors of microsatellite mutuality. However, the interaction between transcription and genomic context with microsatellite mutuality was not significant. We also included mutagenesis index score (MIS) and mutant fitness score (MFS) – measures of gene essentiality from a piggyBac insertion study – from Zhang et al 2018 [29] in the full predictive model (S9 Table), but detected no significant effects.

### Do microsatellite mutation impact phenotype?

Microsatellite length changes are of particular interest if they impact phenotypes. Of the 106 indels observed (98 microsatellites, 2 minisatellites and 6 indels not associated with repeats), 33 are found in promotors, 29 in introns, 25 in intergenic regions and 19 within coding sequences. We examined two metrics for gene essentiality − the MIS and MFS scores [29] − for genes with microsatellite mutations found in this study (S6 Table). Ten of 48 genes were scored as essential in a piggyback mutagenesis screen (MIS < 0.2, MFS < −2), of which two contained coding indels and eight contained intronic indels. We suggest that at least some of the mutations observed will impact phenotype. Given that infected people may contain >10^11^ parasites, and an average mutation rate of 3.00 × 10^-7^/asexual cycle, we expect each infected person to contain an average of >30,000 mutations at each microsatellite locus in the parasite genome. If just a fraction of these mutation modify transcription, then microsatellites may be represent a potent source of genetic variation of which selection can act.

### Comparisons of mutation rate with other species

#### SNPs

To be comparable to other organisms, we translated our base substitution rate to per bp per cell division, assuming 5 cell divisions per *P. falciparum* asexual cycle [30]. This yielded an estimate of 6.19 × 10^-11^ per site per cell division. The per base pair per cell division mutation rates are 3.28 × 10^-10^ for bacteria *Bacillus subtilis* [31], 3.3 × 10^-10^ for yeast *Saccharomyces cerevisiae* [32], 3.2 × 10^-10^ for *Caenorhabditis elegans* [33], 1.5 × 10^-10^ for *Drosophila melanogaster* [34], and 1.0 × 10^-10^ for humans [33, 35, 36]. Hence the estimated *P. falciparum* base mutation rate is comparable to other organisms.

#### Microsatellites

The mutation rate at a microsatellite loci varies according the number of repeated units, length (in base pairs), and the repeated motif [32, 37, 38]. The mutation rates range from 10^-8^-10^-3^ per locus per cell divisions at different microsatellite catalogues for different organisms [37, 38]. The strongest driver of microsatellite mutation is the array length: mutation rate increases as the microsatellite repeat number increases (Fig 8). The mutation rate for *Plasmodium* range from 1.46 (SE: 0.49) × 10^-8^ to 2.11 (0.87) × 10^-7^ per locus per cell division at different motif lengths (1-6 bp), and range from 2.07 (0.86) × 10^-8^ to 1.23 (0.36) × 10^-7^ at different repeat number (3-5 repeats to 20-25 repeats). As all the previous studies focus on mutations with di-, tri, and/or tetra-nucleotide microsatellite mutations, to be comparable, we re-calculated the total mutation rate of di-, tri, and tetra microsatellites at per locus per cell division scale in *P. falciparum* (Fig 8). The overall mutation rates are lower than those reported in human [39–41], *C. elegans* [37] and *S. cerevisiae* [40]. They are also lower than *Dictyostelium discoideum* [42] and *D. melanogaster* [43–45], which have unusually low microsatellite mutation rates. Longer microsatellites have a lower genotyping rate with short-read NGS based methods. To remove this possible bias, we adjusted the *P. falciparum* mutation rate by the average genotype rate for microsatellite repeats of different length (Fig 8). This adjustment increase the microsatellite mutation rate for *P. falciparum* to similar level as *D. discoideum* and *D. melanogaster*. We conclude that the *P. falciparum* microsatellite mutation rate is at the low end of the spectrum compared to other species.

**Fig 8.**
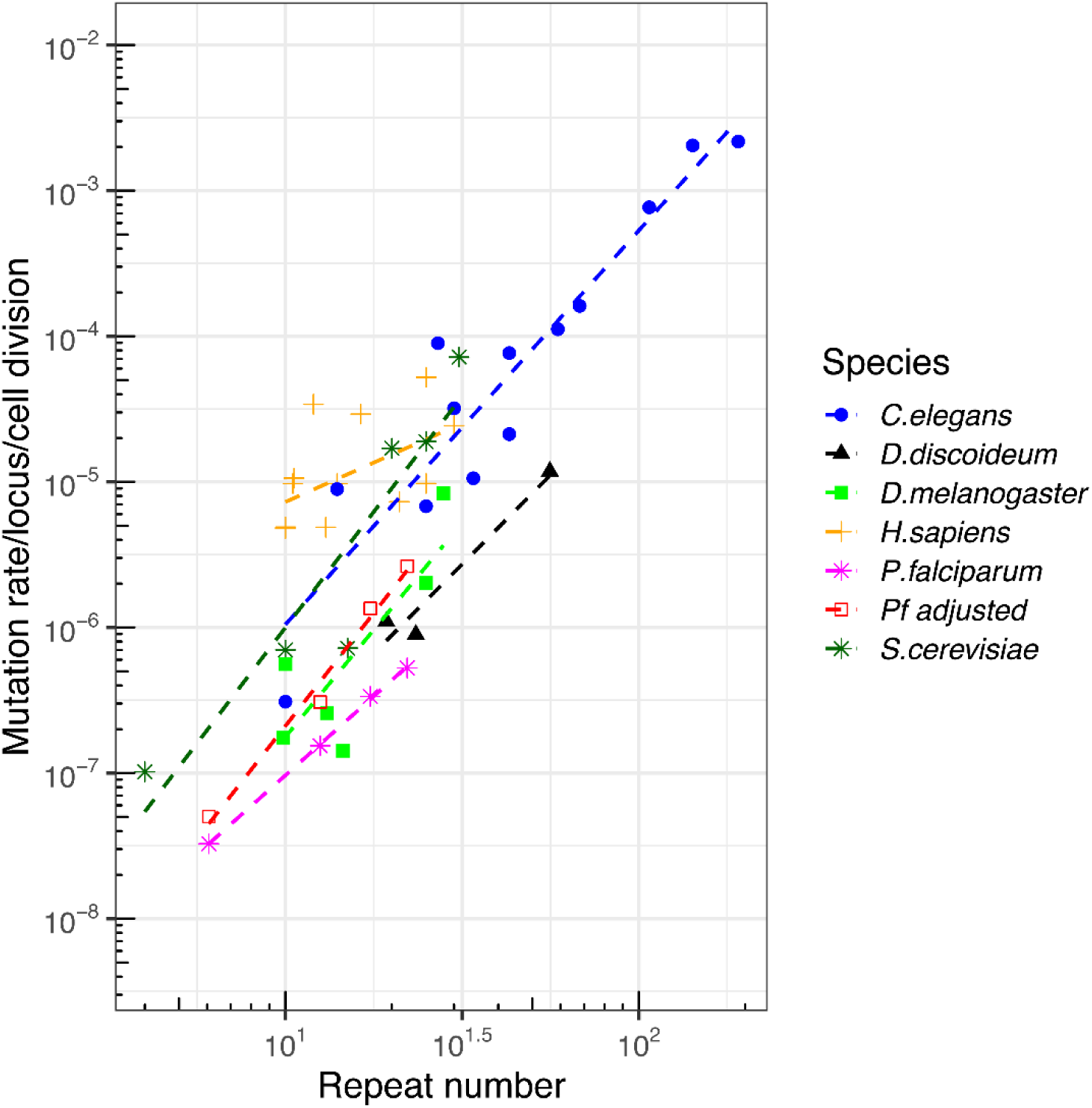
Comparison of mutation rate with other species. We compared published data sets (see text for details) comparing array length and mutation rate across several species. Two lines are provided for our *P. falciparum* dataset, showing unadjusted mutation rates, and mutation rates adjusted by genotyping rate (large microsatellite sequences have lower genotyping rate using short read sequence). *P. falciparum* microsatellite mutation rates are at the low end of the spectrum seen in other organisms.

## Materials and Methods

### Mutation-accumulation process

We generated thirty-one independent mutation-accumulation (MA) lines initiated from a single founder colony of *Plasmodium falciparum* 3D7 (Fig 1). We cultured all parasite lines in complete media at 2% hematocrit and 37°C, with chamber gassed daily and media changed every other day [46]. After every 10.5 parasite cycles, we performed a bottlenecking process to ensure that mutation accumulation is in an effective neutral fashion. Each bottleneck process started with a clonal dilution step; parasites were ultra-diluted into a 96-well plate, to reach a theoretical concentration of 0.25 parasites per well, supplemented with 200 ul of complete media and 2% hematocrit. We identified positive wells with SYBR green I fluorescence assay and confirmed microscopically after 14-21 days post clonal dilution. We transferred the confirmed positive well into a 10 ml flask and cultured for about another week, or until the parasitaemia reached 2%. We then collected the parasites for DNA extraction or cryopreservation, and started another round of bottlenecking. If there was no parasite colony surviving after a particular bottleneck, we would bring up the cryopreserved parasites from the previous bottleneck and again restart another round of parasite dilution and culture. We maintained the MA lines for 114-267 days (S1 Table).

### Whole-genome sequencing and alignment

We extracted DNA with Qiagen DNA Mini kit, sheared about 1.5 µg DNA to ~180bp with Covaris S-series sonicator, and we prepared the sequence libraries with NEBnext library preparation. To reduce the sequencing biases of extremely AT-rich genome, we changed the DNA polymerase to Kapa HiFi DNA polymerase in the PCR enrichment step [47]. We genome sequenced all MA lines and 3D7 progenitor using Illumina Hiseq 2500 platform as described before [48]. Whole-genome sequencing reads for each libraries were individually mapped against the *P. falciparum* 3D7 reference genome (PlasmoDB,http://plasmodb.org/common/downloads/release-32/Pfalciparum3D7/) using alignment algorithm BWA mem [49] under the default parameters. The result pileup files were further converted to SAM format, sorted to BAM format, and deduplicated using Picard tools v2.0.1 (http://broadinstitute.github.io/picard/). We sequenced each sample twice as technical replicates, by equally dividing a single DNA extraction and performing two independent library preparation and Illumina sequencing.

### Genotype calling

We called SNPs and small indels using three methods, including haplotype-based calling method-freeBayes [50], a local *de-novo* assembly-based calling method-GATK HaplotypeCaller [51], and a method designed for genotyping short tandem repeats-HipSTR [15]. To reduce false positives, we excluded genotype callings at variable regions (subtelomeric repeats, hypervariable regions, and centromeres) of the *P. falciparum* genome [2] for all three methods. We applied post-call filtering to each callset as described below (S5 Table).

### Standard filtering parameters

<u>FreeBayes</u>: we first filtered the input aligned reads by requiring a mapping quality ≥30 and base quality ≥20. We excluded reads that didn’t fully span the variable window, when genotyped haplotypes comprised multi-nucleotide polymorphisms (MNPs) or complex variants, to try to detect mutations across repeats. We then generated a raw variant callset which included SNPs, insertions and deletions, MNPs and complex variants under a haploid model (-ploidy 1). We used hard filters (QUAL/AO >10, SAF >0, SAR > 0, RPL > 0 and RPR > 0) to remove low-quality variants. As freeBayes callset included short haplotypes, we normalized the output to show SNPs and indels, for direct comparison to GATK and HipSTR genotypes.

<u>GATK</u>: we followed the best practice recommendations of Genome Analysis Toolkit GATK v3.7, and made slight adaptations for *P. falciparum*. Briefly, base quality score was recalled based on a set of verified known variants [2]. Variants were first called independently for each MA line using HaplotypeCaller and then merged by GenotypeGVCFs, following default parameters but with – sample_ploidy 1. We applied filters to the original GATK genotypes using quality criteria QD > 2, FS < 60, MQ > 40, SOR < 3, GQ > 50 and DP ≥ 3 for SNPs, and QD > 2.0, FS < 200, SOR < 10; GQ > 50 and DP ≥ 3 for indels. As the variant callsets from MA lines are too small, we didn’t perform recalibrate variant quality scores (VQSR) here.

<u>HipSTR</u>: we initially identified microsatellites with at least three repeats, repeat length ranging from 10bp-1000bp, repeat motif size of 1-9bp from the *P. falciparum* 3D7 genome using the *mreps* program [52]. We only genotyped microsatellites located in the core genome for accuracy, and microsatellites shorter than 70bp as genotyping longer microsatellites requires sequence reads longer than 100bp [15]. We performed microsatellite genotyping with HipSTR under the default parameters [15]. Genotypes with any of the following characteristics-< two spanning reads, posterior < 90%, > 15% of reads with a flanking indel, or >15% of reads with a stutter artifact-were excluded from this callset.

### Stringent filtering parameters

Accurate determination of mutation rate requires stringent filtering to remove false positive genotype calls. We used genotype concordance between the two independent sequencing runs for each sample to establish stringent filtering parameters to optimize call accuracy. We used the following statistic to calculate concordance:

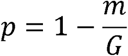

where *p* is the concordance between two independent sequencing runs for an MA line, m is the number of genotype that were discordant in the two runs, and G is the total number of single nucleotide polymorphisms or microsatellite genotyped by both runs.

We first compared genotypes from two separate sequence runs, using callsets generated and filtered as described above (Fig 2, standard filter method). To further remove false positives, we further optimized the filter methods using two parameters, purity and diffPL genotype score. Purity is the percentage of reads that support the current genotype, while diffPL is the difference in likelihood between the reported and next best genotypes. We use conflict calls from two runs as false positive controls to identify appropriate threshold values for stringent filtering of callsets (S4 Fig). We filtered out genotype calls with purity < 0.8, and removed loci where > 50% samples examined failed the purity test. We applied the diffPL filter according to the threshold appropriate for each method (S4 Table). The concordance approached 100% after stringent filter, both SNPs and indels (Fig 2, stringent filter method).

We then collapsed the filtered genotypes from the two sequence runs for each sample by merging identical calls, removing discordant calls and adding calls sequenced from only one run, and combined result from different methods to make the final callset. As a final check, we visually inspected this final genotypes with Integrative Genomics Viewer (IGV, https://software.broadinstitute.org/software/igv/), and manually removed possible errors (S5 Fig).

### Consensus and progenitor-based approach for variant discovery

We used a consensus approach to identify putative mutations in which each individual MA line or progenitor is compared with the consensus genotype of all the remaining lines. This approach identifies variants from large number of samples with low variance in coverage and is robust against sequencing or alignment errors in the reference genome [31, 33, 40, 53, 54]. We employed the consensus approach for our *Plasmodium* dataset with several adjustments: The overall consensus base call is identified as the genotype with the maximum frequency through all the MA lines. We compared the overall consensus with genotypes of progenitor 3D7, and didn’t find any conflict. The individual consensus for each line is compared against the overall consensus. If the line-specific consensus has a call that differs from the overall consensus, and at least two other lines contained enough reads to be used in the comparison, the site was designated as a putative mutation for the discordant line.

### Mutation rate estimates

To calculate the base pair mutation rate per asexual cycle for the MA lines, we used the following equation:

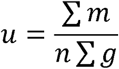

Where *u* is the mutation rate (per base per asexual generation [48 h]), *m* is the number of observed mutations for each MA line, *n* is the number of nucleotide sites analyzed (2.078 × 10^7^ bp for core genome), and *g* is the number of asexual generations for each MA line. We estimated the ninety-five percent confidence intervals and standard error for proportions of polymorphic loci ([∑m]/*n*) by bootstrapping across loci with 1,000 replicates. We counted the number of days from when the progenitor 3D7 was cloned to the final cloning step for each of the MA lines. We then divided this number by the length of an asexual cycle (48h) to determine the total number of asexual cycles for each MA line. We assumed 5 mitotic divisions per 48 h asexual cycle, because a single invading merozoite gives rise to up to 32 (2^5^) daughter merozoites during this time.

To calculate base pair mutation rate for different substitution type, we divided the mutations according to type A:T->G:C, G:C->A:T, G:C->T:A, G:C->C:G, A:T->C:G and A:T->T:A. The number of A/T and G/C nucleotides examined in the core genome were 16,898,216 bp and 3,883,857 bp. The size for coding, promoter (1kb up-stream of start codon ATG), intron and intergenic region were 11,580,293, 4,356,657, 1,407,648 and 3,437,475 bp, respectively in the core genome. The average GC content of these regions are: CDS 23.13%, intergentic region 13.19%, intron 13.93%, and promoters 12.77%. We used the same equation for estimating the mutation rate of microsatellites but with *n* being to the number of microsatellites (123,722) within the core genome. We classified the microsatellite loci by repeat length, repeat motif size and genomic locations to compare the impact of these features on mutation rates (S2-3 Tables).

### Data access

Raw sequencing data have been submitted to the NABI Sequence Read Archive (SRA, https://www.ncbi.nlm.nih.gov/sra) under the project number of PRJNA521718.

## Acknowledgment

We thank National Institutes for Health grant R37 AI048071 (TJCA). This work was conducted in facilities constructed with support from Research Facilities Improvement Program grant C06 RR013556 from the National Center for Research Resources.

## Supporting information

**S1 Fig. Composition of microsatellites in the core genome.** (A) Repeat length, (B) repeat number, (C) representation of microsatellites with different motif sizes in the *P. falciparum* genome. In total there are 123,834 microsatellite tracts in the 3D7 core genome and (D) Distribution of repeat length among microsatellites with different motifs.

**S2 Fig. Composition of the microsatellite within different genomic locations.** (A) Microsatellites are abundant in all regions. (B) Distribution of microsatellites among genome regions. Trinucleotide repeats are enriched in coding regions. (C) Density of microsatellites in different genome regions. Microsatellites are depleted in coding regions. (D-F) repeat number of total, dinucleotide and trinucleotide microsatellite at different genomic regions.

**S3 Fig. Composition of minisatellites repeats in coding and non-coding sequences.** Minisatellites with repeat length divisible by 3 are especially abundant in coding regions.

**S4 Fig. Using conflict calls from two runs as false positives to find the appropriate threshold for stringent filter.** Red dot indicate where the two sequencing runs have different genotype calls; grey dot indicate the same genotype calls; the size of dots show the number of observations.

**S5 Fig.** Visual inspection of putative mutations by IGV plot.

**S6 Fig.** An example for indels with homology sequences.

**S7 Fig. Relationship between mutation rate and repeat array size for dinucleotide microsatellites.** The data fit an exponential better than a linear model (*P* = 0.03). The majority of dinucleotide microsatellites (43,490/43,903, 99.05%) were AT repeats.

**S1 Table.** MA lines and sequence summary.

**S2 Table.** *Plasmodium falciparum* microsatellites information. S3 Table. Composition of the core genome microsatellite.

**S4 Table.** Information of minisatellites.

**S5 Table.** Parameters of standard and stringent filter methods.

**S6 Table.** Mutation number, reproducibility and Concordance under different filter methods.

**S7 Table.** Base substitutions.

**S8 Table.** Indels.

**S9 Table.** Results from the generalized linear model for predicting microsatellite mutability.

## Reference

1. Gardner MJ, Hall N, Fung E, White O, Berriman M, Hyman RW, et al. Genome sequence of the human malaria parasite Plasmodium falciparum. Nature. 2002;419(6906):498.

2. Miles A, Iqbal Z, Vauterin P, Pearson R, Campino S, Theron M, et al. Indels, structural variation, and recombination drive genomic diversity in Plasmodium falciparum. Genome research. 2016;26(9):1288–99.

3. Consortium IHGS. Initial sequencing and analysis of the human genome. Nature. 2001;409(6822):860.

4. Subramanian S, Kumar S. Neutral substitutions occur at a faster rate in exons than in noncoding DNA in primate genomes. Genome research. 2003;13(5):838–44.

5. Anderson TJ, Haubold B, Williams JT, Estrada-Franco § JG, Richardson L, Mollinedo R, et al. Microsatellite markers reveal a spectrum of population structures in the malaria parasite Plasmodium falciparum. Molecular biology and evolution. 2000;17(10):1467–82.

6. Bagshaw AT. Functional mechanisms of microsatellite DNA in eukaryotic genomes. Genome biology and evolution. 2017;9(9):2428–43.

7. Gymrek M. A genomic view of short tandem repeats. Current opinion in genetics & development. 2017;44:9–16.

8. Groh M, Silva LM, Gromak N. Mechanisms of transcriptional dysregulation in repeat expansion disorders. Portland Press Limited; 2014.

9. Lango-Scholey L, Aidley J, Woodacre A, Jones MA, Bayliss CD. High throughput method for analysis of repeat number for 28 phase variable loci of Campylobacter jejuni strain NCTC 11168. PloS one. 2016;11(7):e0159634.

10. Aidley J, Wanford JJ, Green LR, Sheppard SK, Bayliss CD. PhasomeIt: an ‘omics’ approach to cataloguing the potential breadth of phase variation in the genus Campylobacter. Microbial Genomics. 2018.

11. Siena E, D’Aurizio R, Riley D, Tettelin H, Guidotti S, Torricelli G, et al. In-silico prediction and deep-DNA sequencing validation indicate phase variation in 115 Neisseria meningitidis genes. BMC genomics. 2016;17(1):843.

12. Bopp SE, Manary MJ, Bright A T, Johnston GL, Dharia N V, Luna FL, et al. M itotic evolution of Plasmodium falciparum shows a stable core genome but recombination in antigen families. PLoS genetics. 2013;9(2):e1003293.

13. Claessens A, Hamilton WL, Kekre M, Otto TD, Faizullabhoy A, Rayner JC, et al. Generation of antigenic diversity in Plasmodium falciparum by structured rearrangement of Var genes during mitosis. PLoS genetics. 2014;10(12):e1004812.

14. Hamilton WL, Claessens A, Otto TD, Kekre M, Fairhurst RM, Rayner JC, et al. Extreme mutation bias and high AT content in Plasmodium falciparum. Nucleic acids research. 2016;45(4):1889–901.

15. Willems T, Zielinski D, Yuan J, Gordon A, Gymrek M, Erlich Y. Genome-wide profiling of heritable and de novo STR variations. Nature methods. 2017;14(6):590.

16. Mills RE, Luttig CT, Larkins CE, Beauchamp A, Tsui C, Pittard WS, et al. An initial map of insertion and deletion (INDEL) variation in the human genome. Genome research. 2006;16(9):1182–90.

17. Garcia-Diaz M, Kunkel TA. Mechanism of a genetic glissando*: structural biology of indel mutations. Trends in biochemical sciences. 2006;31(4):206–14.

18. Klintschar M, Dauber EM, Ricci U, Cerri N, Immel UD, Kleiber M, et al. Haplotype studies support slippage as the mechanism of germline mutations in short tandem repeats. Electrophoresis. 2004;25(20):3344–8.

19. Forster P, Hohoff C, Dunkelmann B, Schürenkamp M, Pfeiffer H, Neuhuber F, et al. Elevated germline mutation rate in teenage fathers. Proc R Soc B. 2015;282(1803):20142898.

20. Sallmyr A, Tomkinson AE. Repair of DNA double-strand breaks by mammalian alternative end-joining pathways. Journal of Biological Chemistry. 2018:jbc. TM117. 000375.

21. Kirkman LA, Lawrence EA, Deitsch KW. Malaria parasites utilize both homologous recombination and alternative end joining pathways to maintain genome integrity. Nucleic acids research. 2013;42(1):370–9.

22. Ellegren H. Microsatellites: simple sequences with complex evolution. Nature reviews genetics. 2004;5(6):435.

23. Anderson TJ, Su XZ, Roddam A, Day KP. Complex mutations in a high proportion of microsatellite loci from the protozoan parasite Plasmodium falciparum. Molecular ecology. 2000;9(10):1599–608.

24. Ellegren H. Heterogeneous mutation processes in human microsatellite DNA sequences. Nature genetics. 2000;24(4):400.

25. Harr B, Schlötterer C. Long microsatellite alleles in Drosophila melanogaster have a downward mutation bias and short persistence times, which cause their genome-wide underrepresentation. Genetics. 2000;155(3):1213–20.

26. Groh M, Lufino MM, Wade-Martins R, Gromak N. R-loops associated with triplet repeat expansions promote gene silencing in Friedreich ataxia and fragile X syndrome. PLoS genetics. 2014;10(5):e1004318.

27. Zavodna M, Bagshaw A, Brauning R, Gemmell NJ. The effects of transcription and recombination on mutational dynamics of short tandem repeats. Nucleic acids research. 2017;46(3):1321–30.

28. Chen X, Zhang J. Yeast mutation accumulation experiment supports elevated mutation rates at highly transcribed sites. Proceedings of the National Academy of Sciences. 2014;111(39):E4062–E.

29. Zhang M, Wang C, Otto TD, Oberstaller J, Liao X, Adapa SR, et al. Uncovering the essential genes of the human malaria parasite Plasmodium falciparum by saturation mutagenesis. Science. 2018;360(6388):eaap7847.

30. Arnot DE, Ronander E, Bengtsson DC. The progression of the intra-erythrocytic cell cycle of Plasmodium falciparum and the role of the centriolar plaques in asynchronous mitotic division during schizogony. International journal for parasitology. 2011;41(1):71–80.

31. Sung W, Ackerman MS, Gout J-F, Miller SF, Williams E, Foster PL, et al. Asymmetric context-dependent mutation patterns revealed through mutation–accumulation experiments. Molecular biology and evolution. 2015;32(7):1672–83.

32. Lynch M, Sung W, Morris K, Coffey N, Landry CR, Dopman EB, et al. A genome-wide view of the spectrum of spontaneous mutations in yeast. Proceedings of the National Academy of Sciences. 2008.

33. Denver DR, Dolan PC, Wilhelm LJ, Sung W, Lucas-Lledó JI, Howe DK, et al. A genome-wide view of Caenorhabditis elegans base-substitution mutation processes. Proceedings of the National Academy of Sciences. 2009;106(38):16310–4.

34. Haag-Liautard C, Dorris M, Maside X, Macaskill S, Halligan DL, Charlesworth B, et al. Direct estimation of per nucleotide and genomic deleterious mutation rates in Drosophila. Nature. 2007;445(7123):82.

35. Giannelli F, Anagnostopoulos T, Green P. Mutation rates in humans. II. Sporadic mutation-specific rates and rate of detrimental human mutations inferred from hemophilia B. The American Journal of Human Genetics. 1999;65(6):1580–7.

36. Kondrashov AS. Direct estimates of human per nucleotide mutation rates at 20 loci causing Mendelian diseases. Human mutation. 2003;21(1):12–27.

37. Seyfert AL, Cristescu ME, Frisse L, Schaack S, Thomas WK, Lynch M. The rate and spectrum of microsatellite mutation in Caenorhabditis elegans and Daphnia pulex. Genetics. 2008;178(4):2113–21.

38. Bhargava A, Fuentes F. Mutational dynamics of microsatellites. Molecular biotechnology. 2010;44(3):250–66.

39. Brinkmann B, Klintschar M, Neuhuber F, Hühne J, Rolf B. Mutation rate in human microsatellites: influence of the structure and length of the tandem repeat. The American Journal of Human Genetics. 1998;62(6):1408–15.

40. Lynch M, Sung W, Morris K, Coffey N, Landry CR, Dopman EB, et al. A genome-wide view of the spectrum of spontaneous mutations in yeast. Proceedings of the National Academy of Sciences. 2008;105(27):9272–7.

41. Hohoff C, Dewa K, Sibbing U, Hoppe K, Forster P, Brinkmann B. Y-chromosomal microsatellite mutation rates in a population sample from northwestern Germany. International journal of legal medicine. 2007;121(5):359–63.

42. McConnell R, Middlemist S, Scala C, Strassmann JE, Queller DC. An unusually low microsatellite mutation rate in Dictyostelium discoideum, an organism with unusually abundant microsatellites. Genetics. 2007.

43. Schug MD, Hutter CM, Wetterstrand KA, Gaudette MS, Mackay T, Aquadro CF. The mutation rates of di-, tri-and tetranucleotide repeats in Drosophila melanogaster. Molecular Biology and Evolution. 1998;15(12):1751–60.

44. Schlötterer C, Ritter R, Harr B, Brem G. High mutation rate of a long microsatellite allele in Drosophila melanogaster provides evidence for allele-specific mutation rates. Molecular Biology and Evolution. 1998;15(10):1269–74.

45. VÁzquez JF, PÉrez T, Albornoz J, DomÍnguez A. Estimation of microsatellite mutation rates in Drosophila melanogaster. Genetics Research. 2000;76(3):323–6.

46. Trager W, Jenson JB. Cultivation of malarial parasites. Nature. 1978;273(5664):621–2.

47. Oyola SO, Otto TD, Gu Y, Maslen G, Manske M, Campino S, et al. Optimizing Illumina next-generation sequencing library preparation for extremely AT-biased genomes. BMC genomics. 2012;13(1):1.

48. Cheeseman IH, McDew-White M, Phyo AP, Sriprawat K, Nosten F, Anderson TJ. Pooled sequencing and rare variant association tests for identifying the determinants of emerging drug resistance in malaria parasites. Molecular biology and evolution. 2015;32(4):1080–90.

49. Li H. Aligning sequence reads, clone sequences and assembly contigs with BWA-MEM. arXiv preprint arXiv:13033997. 2013.

50. Garrison E, Marth G. Haplotype-based variant detection from short-read sequencing. arXiv preprint arXiv:12073907. 2012.

51. Poplin R, Ruano-Rubio V, DePristo MA, Fennell TJ, Carneiro MO, Van der Auwera GA, et al. Scaling accurate genetic variant discovery to tens of thousands of samples. BioRxiv. 2017:201178.

52. Kolpakov R, Bana G, Kucherov G. mreps: efficient and flexible detection of tandem repeats in DNA. Nucleic acids research. 2003;31(13):3672–8.

53. Ossowski S, Schneeberger K, Lucas-Lledó JI, Warthmann N, Clark RM, Shaw RG, et al. The rate and molecular spectrum of spontaneous mutations in Arabidopsis thaliana. science. 2010;327(5961):92–4.

54. Sun Y, Powell KE, Sung W, Lynch M, Moran MA, Luo H. Spontaneous mutations of a model heterotrophic marine bacterium. The ISME Journal. 2017;11(7):1713–8.

