## Supplemental Figures for "Mode and tempo of microsatellite length change in a malaria parasite mutation accumulation experiment"

### Supplementary Information

**S1 Fig**. **Composition of microsatellites in the core genome.** (A) Repeat length, (B) repeat number, (C) representation of microsatellites with different motif sizes in the *P. falciparum* genome. In total there are 123,834 microsatellite tracts in the 3D7 core genome and (D) Distribution of repeat length among microsatellites with different motifs.

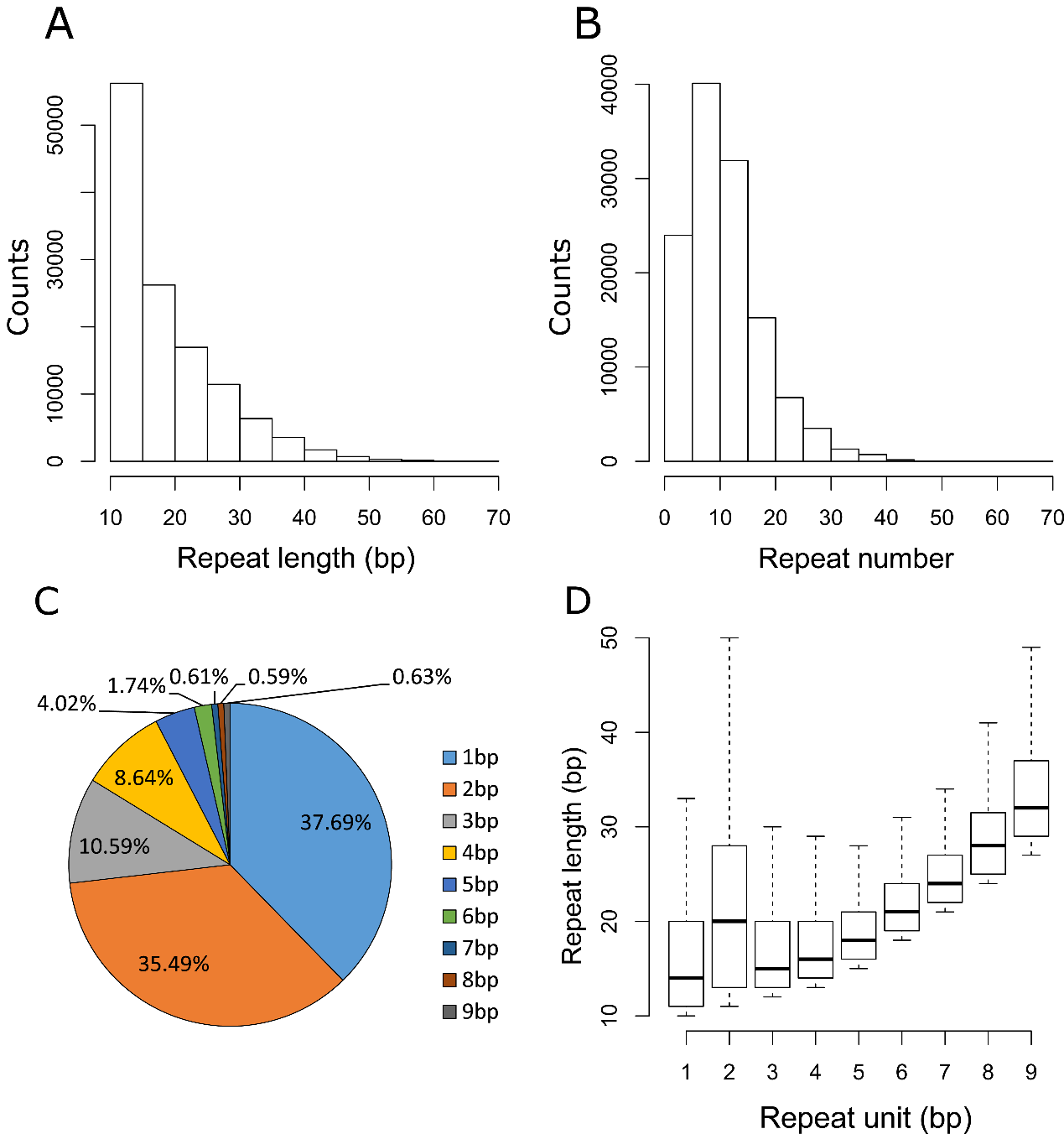

**S2 Fig. Composition of the microsatellite within different genomic locations.** (A) Microsatellites are abundant in all regions. (B) Distribution of microsatellites among genome regions. Trinucleotide repeats are enriched in coding regions. (C) Density of microsatellites in different genome regions. Microsatellites are depleted in coding regions. (D-F) repeat number of total, dinucleotide and trinucleotide microsatellite at different genomic regions.

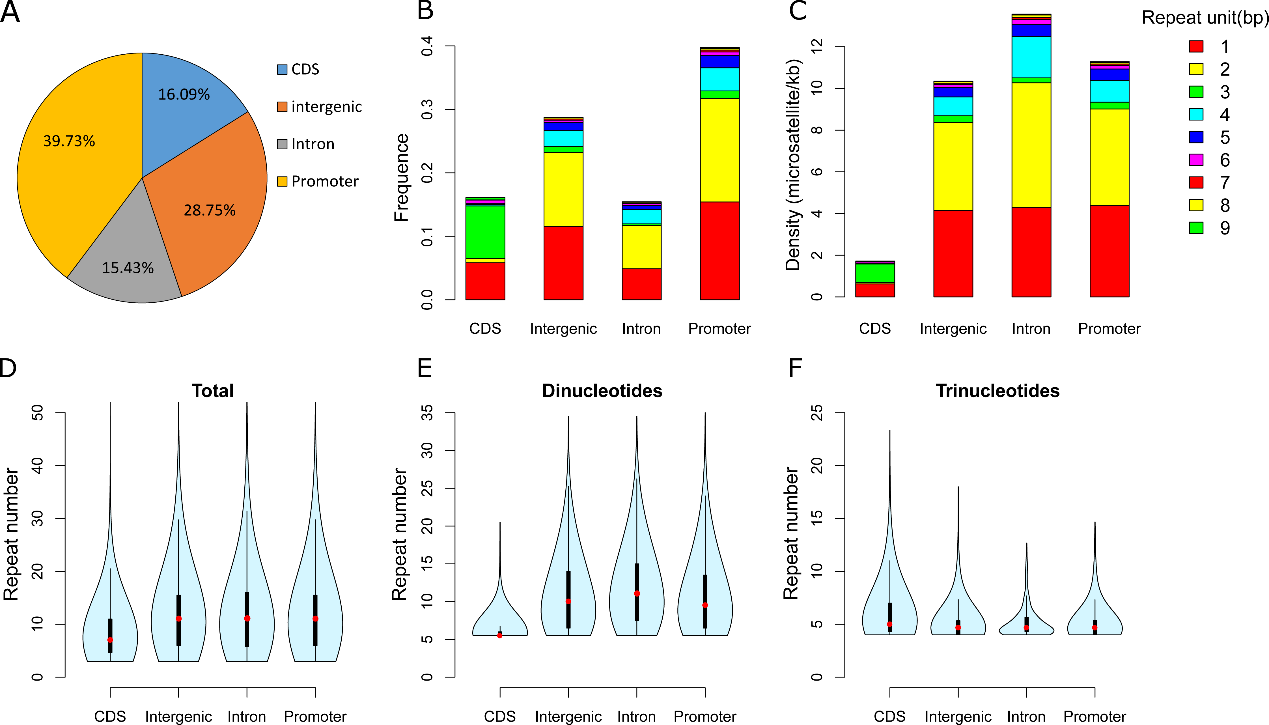

**S3 Fig. Composition of minisatellites repeats in coding and non-coding sequences.** Minisatellites with repeat length divisible by 3 are especially abundant in coding regions.

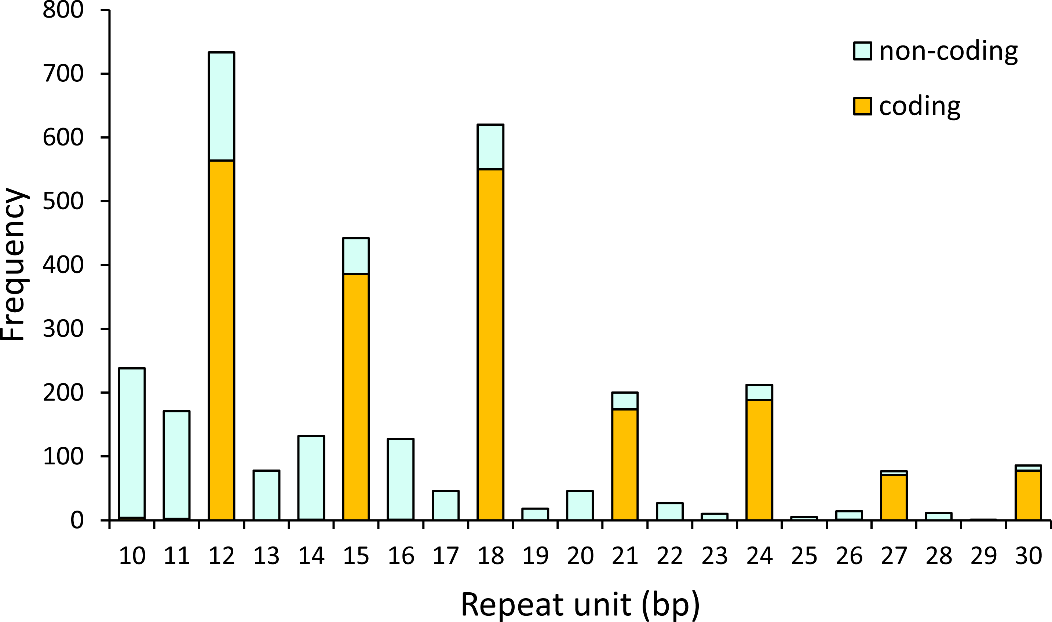

**S4 Fig. Using conflict calls from two runs as false positives to find the appropriate threshold for stringent filter.** Red dot indicate where the two sequencing runs have different genotype calls; grey dot indicate the same genotype calls; the size of dots show the number of observations.

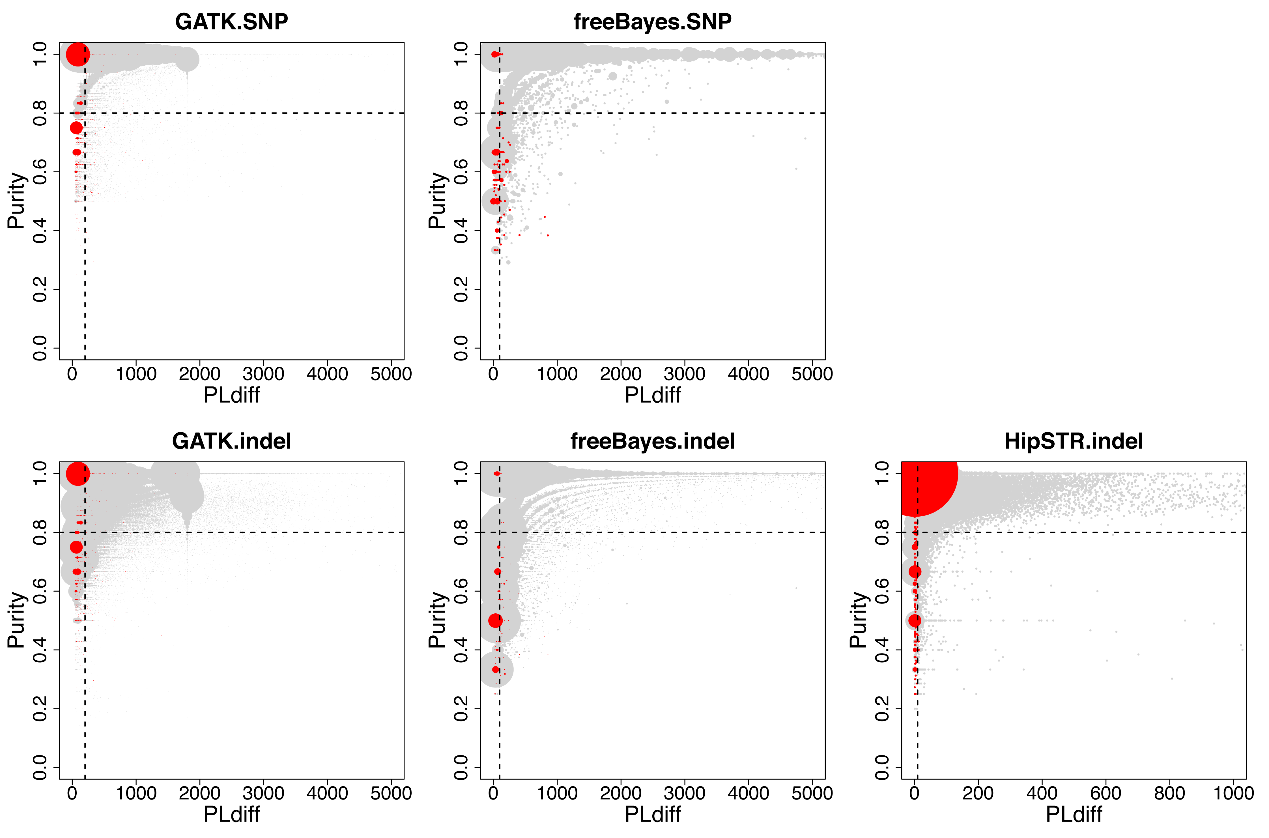

**S5 Fig. Visual inspection of putative mutations by IGV plot.**

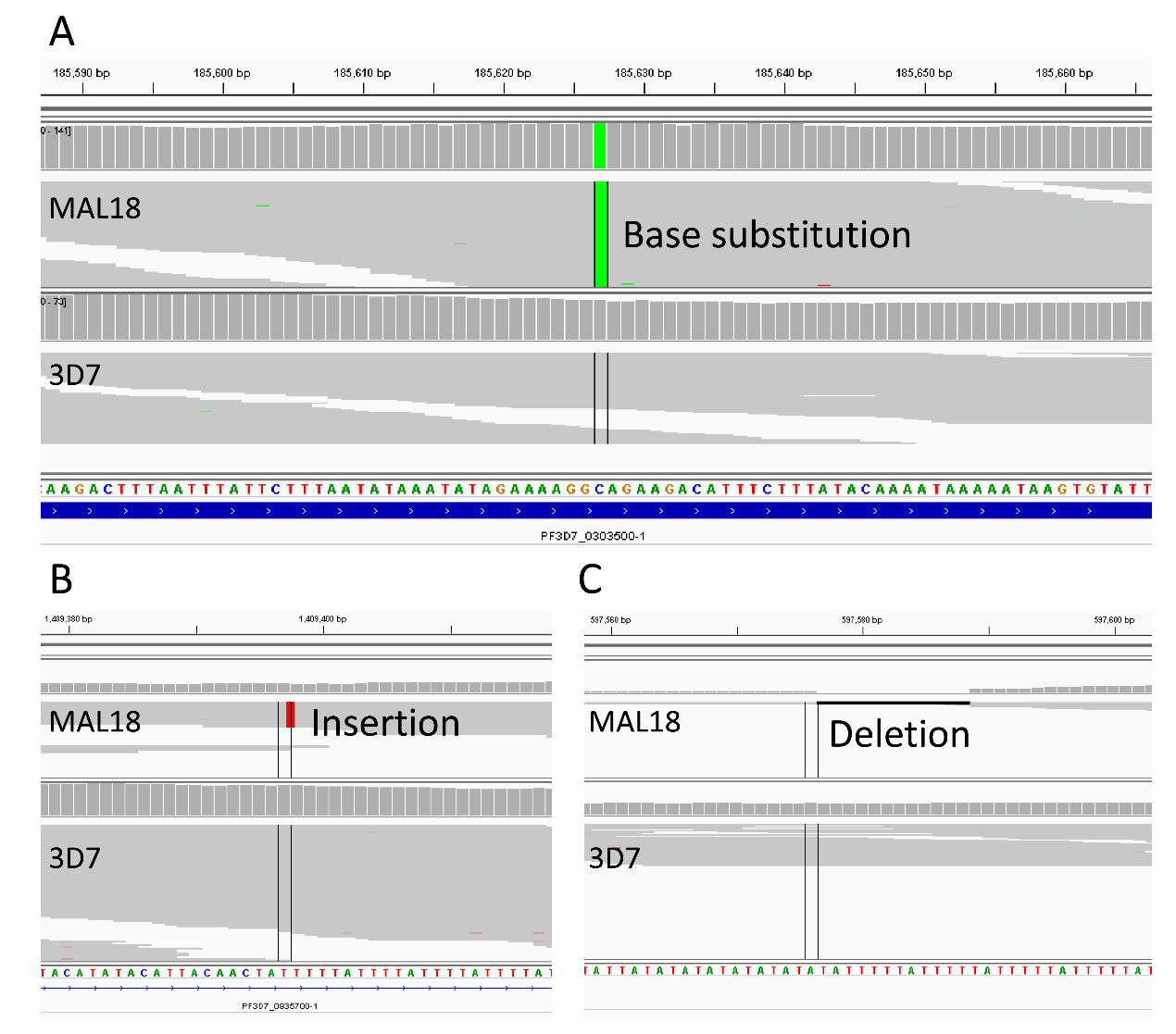

**S6 Fig. An example for indels with homology sequences.**

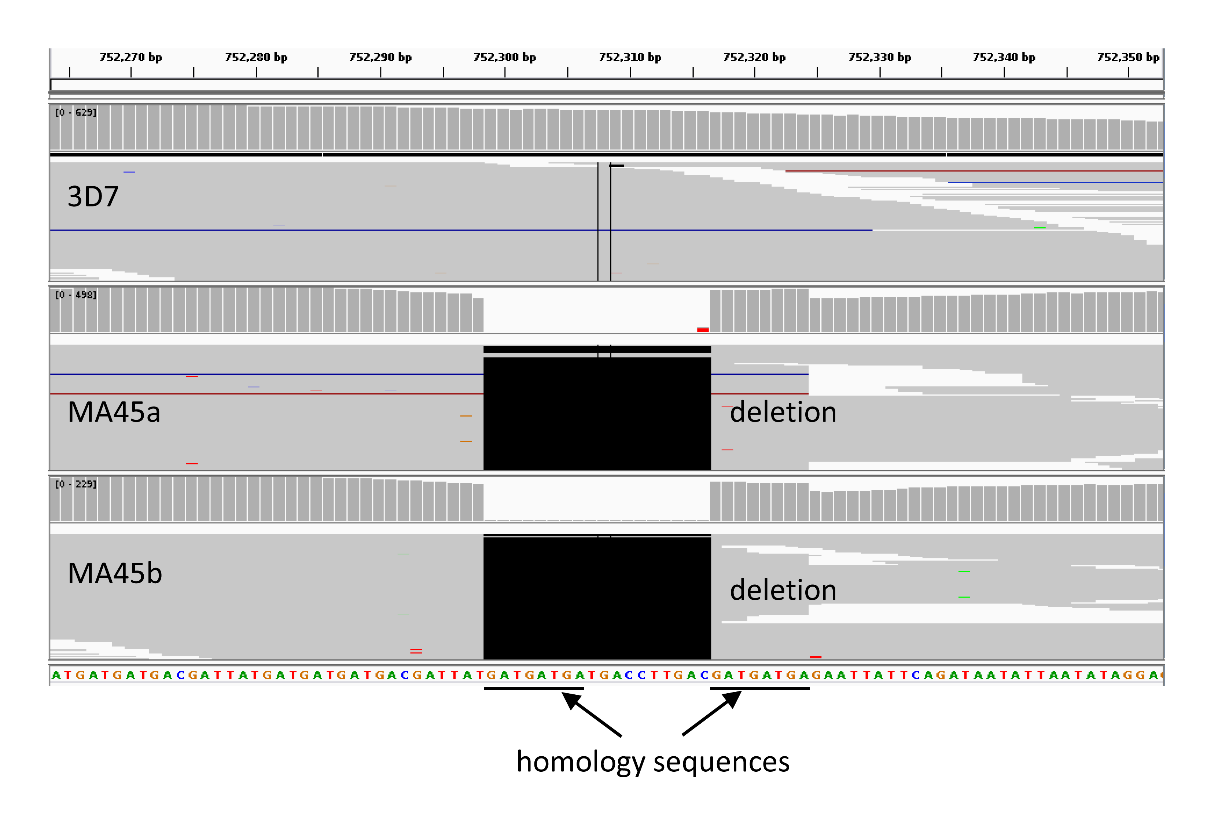

**S7 Fig. Relationship between mutation rate and repeat array size for dinucleotide microsatellites.** The data fit an exponential better than a linear model (*P* = 0.03). The majority of dinucleotide microsatellites (43,490/43,903, 99.05%) were AT repeats.

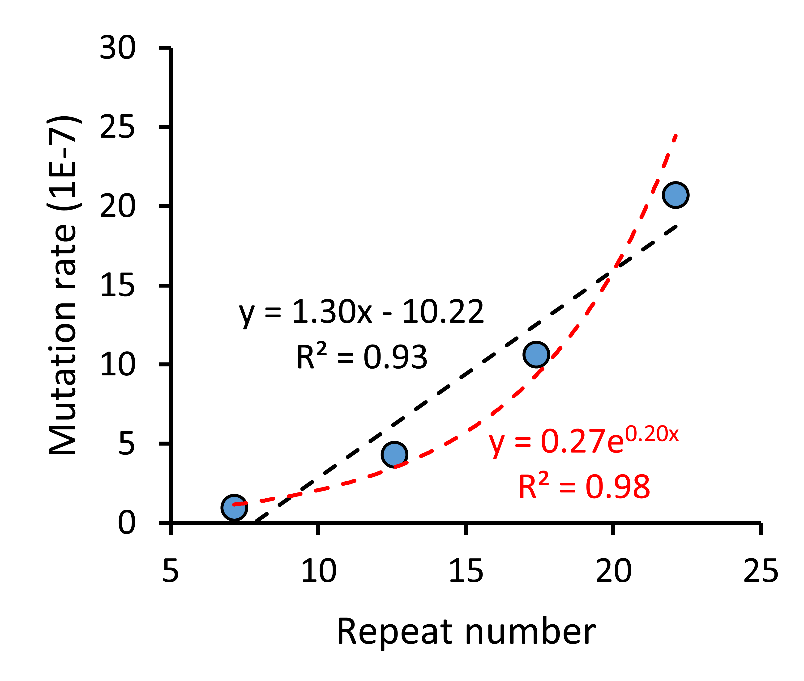
